# Chromatin configuration affects the dynamics and distribution of a transiently interacting protein

**DOI:** 10.1101/246231

**Authors:** A. Amitai

## Abstract

We present a theoretical study of the interaction between a protein (diffusing particle) with chromatin (polymer chain). Each monomer is a trap where a particle can transiently bind. We derive novel formulas for the transition rate between monomer sites, given a specific polymer configuration, and find that a particle is likely to rapidly rebind many times to its release site, before moving to another. The reattachment probability is larger when the local density around the release site is smaller. Interestingly, for an equilibrated polymer, the transition probability decays as a power-law for close monomer-to-monomer distances and reaches an asymptotic value for faraway ones. By computing the transition rate between monomers, we show that the problem of facilitated search by a protein can be mapped to a continuous time Markov chain, which we solve. Our findings suggest that proteins may be locally trapped for a time much longer than their dissociation time, while their overall motion is ergodic. Our results are corroborated by Brownian simulations.

---

The interaction of proteins with chromatin regulates many cellular functions. Most DNA-binding proteins interact both non-specifically and transiently [1] with many chromatin sites as well as specifically and more stably with cognate binding sites. These interactions and chromatin structure are important in governing protein dynamics [2, 3]. However, the effect of these transient interactions on protein motion and distribution has yet been shown from a first principle.

Some aspects of protein interactions with DNA have been studied in the context of the search of a gene promoter site by a transcription factor (TF) [4]. It was first noted [5] that the search for a promoter site by a TF would be faster if it involves 3d excursions, as well as sliding of the protein along DNA [6], as was shown in prokaryotes [7]. These different types of motion were observed experimentally, leading to massive interest in models of facilitated diffusion [8–15], and in the impact of a regulating site’s position on transcription [16, 17]. In current microscopy experiments, it is impossible to examine the search process to its fullest [18]. Thus, we concentrate here on modeling the experimentally observed dynamics of the protein as a diffusing and interacting particle.

We will show that proteins are likely to stay in the proximity of a site for a time much longer than their dissociation time from DNA due to reattachment. Moreover, we find that reattachment depends on the local density around the release site and that the precise configuration of the polymer impacts the interacting particle’s dynamics. We further find the rates associated with the transition between different monomer sites. Finally, we show that the process as a whole is ergodic; it has no long-time power law distribution of the residence time at a site as has been previously suggested [20].

We consider a point particle (protein) placed at a distance *a* from monomer *n* (locus), part of a long flexible polymer [21, 22] (chromatin) (Fig. 1a). The particle diffuses until encountering monomer *l*. Absorption occurs at capture radius *∊* < *a* from *l* (Fig. 1b), where it remains bound for characteristic time *T*. Upon dissociation, the particle is placed at a distance *a* from monomer *l* position (Fig. 1b), with a uniform angular distribution and starts diffusing again. While we postulate here two radii: release (*a*) and capture (*∊*), the effective behavior is equivalent to a model with one release radius, and a partially reflecting boundary condition on it. In a partially reflecting model, the particle has a probability to re-absorb immediately after release. Small re-absorption probability corresponds to *∊* ≪ *a*. Thus, when the release and capture radii are comparable, the particle spends much time around one monomer cluster, then jumps to another, only to come back to the first (Fig. 1c).

**Fig. 1.**
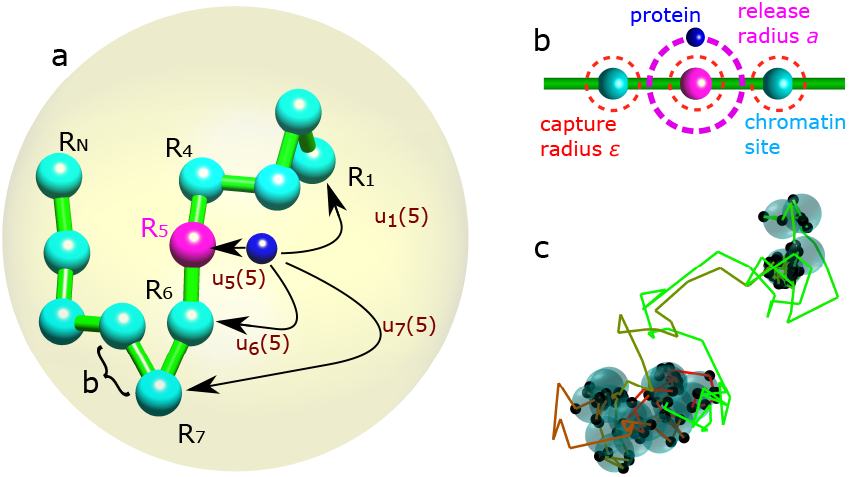
Transition between monomer sites. (a) A particle (dark blue) representing a protein interacts with a monomer site (cyan) that serves as a chromatin locus. It detaches from an initial monomer *n* (magenta) and reattaches to site *l* with probability *u_l_*(*n*) without touching other sites along the way. The polymer is inside a confining domain (yellow) of radius *A*. *b* is the mean square displacement of a bond. (b) Theparticleisreleasedatdistance *a* from the initial site, with uniform angular distribution. It attaches to a site once it is at distance *∊* from it. We depict chromatin as a coarse-grained chain of beads, each of characteristic size *b* representing 3.2kb and of size 30nm [19]. Hence, for *∊* = 0.03*b* = 0.9nm and *a* = 0.3*b* = 9nm, the release and capture radii are of the order of the size of a protein and the interaction distance. (c) Trajectory of the particle interacting with the monomer sites (cyan). The trajectory color changes with time from red to green. The black dots are attachment points at the monomer site (*a* = 0.5*b*, *∊* = 0.49*b, N* = 100, *A* = 6*b*).

The probability of the particle to arrive at a certain monomer site before another via 3d diffusion depends on their respective initial distance [23]. The probability *u_l_*(*x*) that a particle starting from *x* arrives at monomer *l* before encountering any other monomer (see SM) is 
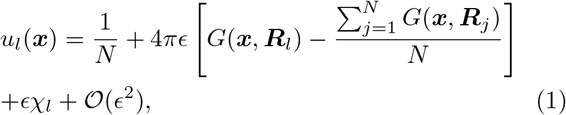
where *N* is the polymer length, ***R_i_*** is the position of monomer *i, G(x, y)* is the Neumann Green’s function of the Laplacian in a sphere of radius *A, χ_l_* is a constant that depends on the monomers’ positions but not on the initial position *x*. In writing eq.(1), we assume that the trapping monomers are well separated. When the *∊* neighborhoods of two monomers merge, the equation can be modified [24].

Proteins move much faster in the nucleus than chromatin. The diffusion coefficient of a chromatin locus can be estimated by inserting a fluorescent tag and following its trajectory [25]. Assuming a coarse-grained model of chromatin, it was found that a locus of size 3*kb* has a diffusion coefficient of about *D* = 10^*−*2^*μm*^2^/sec [22], while proteins move three orders of magnitude faster- For example, 13.5*μm*^2^/sec for the c-Myc protein [26]. We thus assume that the polymer is fixed at an equilibrium configuration inside the domain while the particle is diffusing.

Assuming that the polymer is equilibrated in the domain, we found the transition probability *u_l_*(*n*) that a particle starting in the proximity of site *n* finds site *l* first before encountering other monomers by averaging eq.(1) with the equilibrium distribution of the polymer in bulk (see SM): 
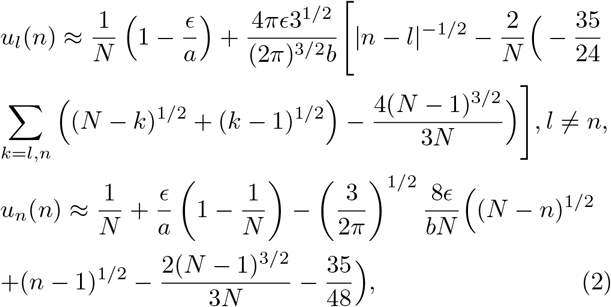
where 2 < *n, l* < *N,b* is the standard deviation of a bond length (Fig. 1a) and we assume *∊* (*N*^1/2^/b + *Ca^−1^*) ≪ 1, with *C* a numerical factor of order 1. The expression for the end monomers is different (see SM sec.4).

While *N* we did not explicitly include a sliding state of the protein along DNA [6], it could be included by modifying the nearest-neighbors transition probability (*u_n−1_*(*n*), *u_n_*_+1_(*n*)) and will not change qualitatively our results.

The reattachment probability *u_n_*(*n*) at the release site is larger than the probability of attaching to far-away sites (Fig. 2a, eq.2), suggesting that once released, the particle is likely to rebind at the same site. For a longer polymer strand, the relative probability between reattachment and attachment to a faraway site is larger (Fig. 2b). This result is contrary to previous studies that assumed that the transition probability and the *k_on_* for rebinding was equal between all monomers [13, 27–33], or related to a Lévy type diffusion of the particle [31].

**Fig. 2.**
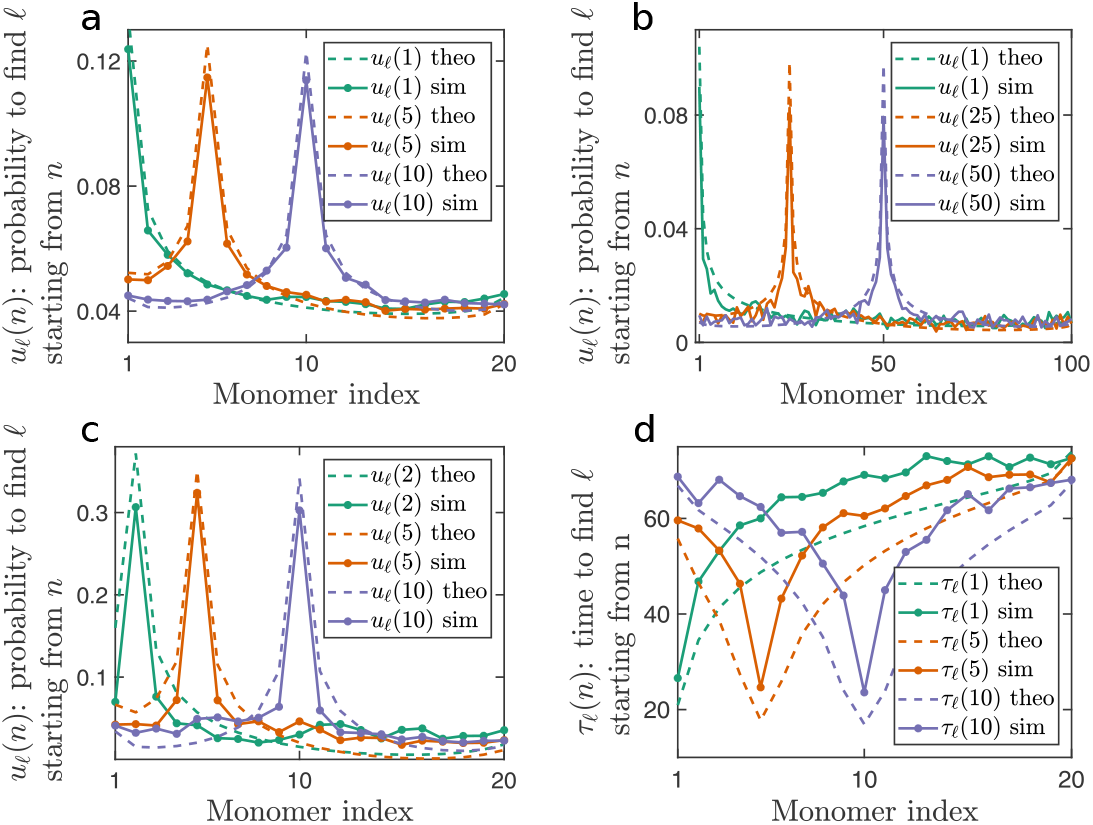
Transition between monomer sites. (**a**) The transition probability starting from site 1 (turquoise), 5 (orange) or 10 (purple). The polymer has 20 monomer sites (*N* = 20), *a* = 0.3*b*, *∊* = 0.03*b, A* = 10*b*. The full lines are the result of Brownian simulations, while the dashed lines are computed using the analytical formula (2). (**b**) The site transition probability when the polymer is longer (*N* = 100). (**c**) The polymer is of length *N* = 20. It is crowded in a small domain *A* = 2*b* and the capture radius is *∊* = 0.12*b*. (**d**) The mean first transition time *τ_l_*(*n*) from site *n* to site *l* without interacting with any other site along the way. The full lines are the result of Brownian simulation and the dashed line is computed using formula (5) with the same parameters as in(a). A single time unit is equal to 1t.u = 10*b*^2^*/D*, with *D* the diffusion coefficient of the particle.

Interestingly, we find that the reattachment probability is minimal for the middle monomer (eq.2). To clarify this effect, we estimated the expected number of monomers in a ball around the release monomer, for different monomers along the chain (see SM Fig. 1). This quantity is akin to the local density around the release site. The monomer density is highest around the middle monomer, which is closest to the polymer center of mass. Thus, the reattachment probability is sensitive to the local density around the release site and not just to the average monomer density in the domain.

Chromosomes’ segregation in the nucleus may be the result of self-avoiding interaction (SAI) between them [34]. To study how SAI between monomers would modify the behavior of *u_l_*(*n*), we performed Brownian simulation (BS) where the monomers interacted through the Lennard-Jones potential [35] in addition to nearest-neighbors spring interaction (see SM). The separation of monomers due to SAI decreases the local monomer density around the release point, leading to a larger reattachment probability *u_n_*(*n*) compared to a phantom chain (see SM Fig. 2a).

The root mean square distance between monomers scales with their distance along the chain 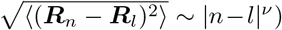. This results in the power-law scaling *u_l_*(*n*) ∼ |*n − l*|^−1/2^ for proximal sites of a phantom polymer (eq.2), for which *ν* = 1/2. To study the behavior for a SA polymer, we computed *u_l_*(*n*) between proximal sites from the BS, averaged over all release sites *n* and fitted it: 〈*u_|l−n|_*(*n*)∼ *A*|*l* − *n*|^−α^ + *C*. We found the fitted exponent to be *α* = 0.59 (see SM Fig. 2b) as would be expected from the rmsd of a SA polymer, for which *ν* = 0.6 is the Flory exponent [36]. When *a* and *∊* are larger, *α* increases (see SM Fig. 2c). This behavior may be explained by studying high order expansions of *u_l_*(*x*) in the parameter *∊*.

Computing *u_l_*(*n*), we assumed that the polymer configuration is not much affected by confinement (the ratio of gyration radius to the domain radius: 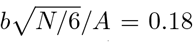 and *R_g_/A* = 0.4 for Fig. 2a and b respectively). Thus, its equilibrium distribution was well approximated by the equilibrium distribution of a flexible chain in bulk. Since one end of the polymer is anchored at the origin of the domain, when the end-to-end distance 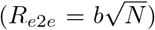 is of the order of the domain radius, the polymer feels the effect of confinement.

In the nucleus, chromosomes are tightly packed and are not in the dilute regime. We thus simulated a polymer in a smaller domain for which *R_g_/A* = 0.91 and *R_e_*_2*e*_/A = 2.23 and computed *u_l_*(*n*) (Fig. 2c). Interestingly, eq.(2) still matches the transition probability even for this moderately crowded polymer. Indeed, when the target monomer *l* is close by, its distance distribution from the release monomer is not affected much by the presence of confinement. At the same time, the transition probability to far away monomers weakly depends on the distance between the release site and the target. Thus, it is not much affected by confinement. The validity of eq.(2) is expected to break for extreme packing (*A* ≪ *R_g_*).

To understand the rates involved in the encounters between proteins and chromatin, we computed the conditional mean first passage time (MFPT) *τ_l_*(*x*) of a particle from position *x* to site *l* without encountering other monomers along the way. The conditional dynamics of the particle obeys the Langevin equation [37] 
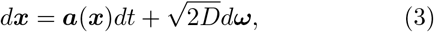
where *dω* is a white Gaussian noise, the drift 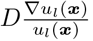, and *u_l_*(*x*) is given by eq.2. Since *u*(*x*) approaches zero when *x* approaches any monomer other than *l* (see SM eq.5), the drift *a*(*x*) will diverge for *x* → *R_i_* (*i* ≠ *l*). Thus, moving according to eq.(3), the particle experiences a drift pushing it away from all monomers except for its final destination *l*.

*τ_l_*(*x*) can be found by solving a boundary value problem (see SM eq.73). It has the approximate solution 
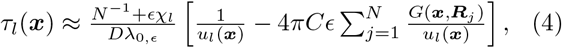

Where *λ*_0,*∊*_ is the eigenvalue of an associated eigenvalue problem (see SM sec. 5).

Using the pre-averaging technique on eq.(4) with the equilibrium polymer configuration, we find an asymptotic formula for the mean conditional transition time starting from monomer *n* 
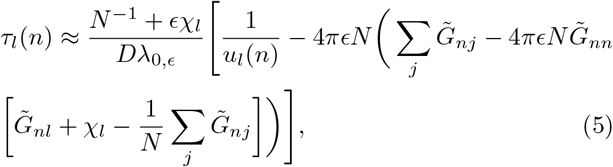
where *G_nj_* is the Green’s function between monomers *n* and *j* positions, averaged over the equilibrium distribution.

Compared to BS, formula (5) matches the simulation, where the difference is up to about 20% of the MFPT (see Fig. 2d). Interestingly, *τ_l_*(*n*) to the release site (*l* = *n*) is much faster than to other sites and the MFPT converges asymptotically when the target monomer is far from the release site (*|l − n|* ≫ 0). When *∊ → a*, the recapture time is faster with respect to transition time to another site. Indeed, when an interacting protein starts from the boundary layer of the release site, the characteristic recapture rate is significantly higher than the travel time to other sites. Hence, we suggest that when a protein is released from a chromatin site, it would quickly rebind to it with high probability.

We find that the addition of SAI reduces *τ_l_*(*n*) compared to a phantom polymer (SM Fig. 2d). The addition of SAI increases the average distance between monomers. Consequently, the eigenvalue *λ*_0,*∊*_ is larger (see SM eq.87), resulting in a smaller MFPT (eq.5). Therefore, the monomers of a SA polymer fill the domain more optimally than those of a phantom polymer, resulting in a rapid capture.

Since the reattachment rate at a site is larger, we expect that when SAI are dominant, proteins will spend a larger fraction of their time bound at monomer sites.

Heterochromatin is considered to be denser than euchromatin [38]. However, the nature of proteins’ interaction with heterochromatin it is still unclear. We would expect *u_n_*(*n*) to be smaller in heterochromatin domains and a protein to “forget” faster its release position. Alternatively, if reattachment occurs with higher probability (*∊ → a*) in heterochromatin, *u_n_*(*n*) will be larger in these domains.

Proteins in the nucleus can either be bound to chromatin, to other nuclear compartments or stochastically moving between association events. We denote the unbound state as *in-transit* from one monomer site to another. We assume that at site *n*, the particle remains bound for a characteristic time *T_n_*, and leaves with a Poissonian rate. We thus formulated the transition between the different states using a continuous time Markov chain (CTMC) (Fig. 3a). We constructed the rate matrix *Q* between bound states and transit-states that depend on *u_l_*(*n*), *τ_l_*(*n*) and *T_n_* (see SM). We used *Q* to find the time evolution of the probability distribution function of the particle

**FIG. 3.**
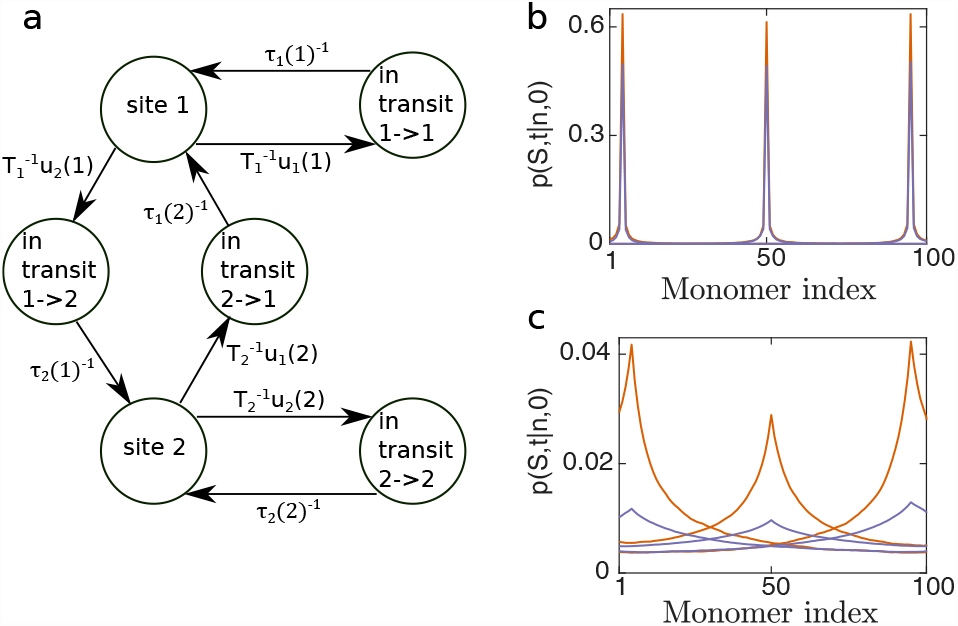
Site-site transition time and particle distribution in the domain. (**a**) Formulating the behavior of a particle (protein) that interacts with the polymer (chromatin), inside a domain (nucleus) as a continuous time Markov chain. We illustrate it for the case of two binding monomer sites. The particle can be bound at site 1 or 2 and is released with Poissonian dissociation rates 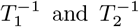 While diffusing to bind to a site, it is “in-transit” and arrives at its destination with probability one and rate *τ_l_*(*n*)^*−1*^. (**b-c**) The monomer is bound for a characteristic time *T* = 1t.u at a site. The probability *p* of the particle to be in a bound state at monomer after 1*T* (b) or 10*T* (c), is computed by solving numerically eq.(6). The monomer start at the *a*-neighborhood of either monomer 2, 50 or 99. *u_l_*(*n*) and *τ_l_*(*n*) were estimated from Brownian simulation: *N* = 100, *a* = 0.5*b, A* = 10*b*, *∊* = 0.49 (orange) or *∊* = 0.3*b* (purple). For *N* = 100 there are 100 bound states and 10^4^ in-transit states.

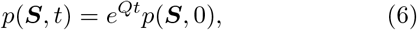

where *S* is the state vector of the CTMC and *p*(*S*, 0) is the initial particle distribution.

We performed BS and estimated numerically *u_l_*(*n*)and *τ_l_*(*n*), taking *T_n_* = *T* (*n* = 1*…N*) and estimated *p*(*S,t*) starting from site *n* using eq.(6). When the reattachment probability is high, the particle remains in proximity to its initial site for a long time (Fig. 3b,c) compared with *T*. Thus, the equilibration time of the protein in the domain is much longer than its disassociation rate from a site (*k_off_* = 1*/T*). When the reattachment probability is smaller, the particle diffuses farther from its initial site (Fig. 3c). Interestingly, the residence probability at the original site is not uniform along the chain. Since the middle monomer resides where monomer density is highest, the reattachment probability *u*_N/2_(*N*/2) is minimal (eq.2) for the middle monomer.

Using the long-time behavior of *p*(*S,t*), we estimated the fraction *f* of bound particles. For high reattachment probability *f*^*ε* = 0.49*b,a* =0.5*b*^ ≈ 0.93, while the rest of the probability is in the unbound (in-transit) states. For smaller reattachment probability *f*^*e* = 0.49*b,a* =0.5*b*^ ≈ 0.6. In the SM we plotted *f* for different values of *∊* (see SM Fig. 3). *f* estimated using eq.6 corresponds to the bound fraction estimated directly with BS.

A naive estimate for the bound fraction, that does not take into account reattachment, can be found through the ratio of the off-rate *k_off_* and on-rate starting from the bulk 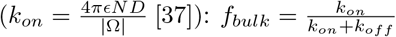. We found that the bulk estimate greatly underestimates the bound fraction found from the BS or using the CTMC formalism (see SM Fig. 3). Thus, starting at the boundary layer of the initial monomer site can impact significantly proteins bound fraction. This may be the origin of the observed non-uniform distribution [39] and protein clusters [40] in the nucleus.

To conclude, there is a finite probability, after each release, that the protein diffuses away to another remote site, given that there is a trajectory between them. Thus, the process is ergodic. When the *∊*-neighborhoods of many traps overlap around the released site, such that the particle cannot find a path out, it will be quenched in this area.

Our findings can explain the long residence time distributions that are observed experimentally [20] without the need for a power law waiting time distribution as assumed in a CTRW model. As we have shown, escaping a binding site involves several dissociation and association events, with different characteristic rates. Hence, the localization time distribution may not appear to have exponential distribution in experiments.

Since the 3d organization of chromatin guides search of TFs through transient interactions, the number of proteins and their interaction strength is not sufficient to understand their collective behavior. To fully model proteins behavior at chromatin loci, one has to study the nature of the local interactions around the site of interest. Based on our model, we can extract directly from microscopy data the interaction parameters of proteins at specific chromatin domains. Thus, we can understand how different proteins “see” chromatin differently.

## AUTHOR CONTRIBUTIONS

A.A. performed the research and wrote the paper.

## ACKNOWLEDGMENTS

The author thanks M. Kardar, D. Holcman, J. Reingruber and A. K. Chakraborty for their helpful discussions and comments. I would like to acknowledge financial support from Massachusetts General Hospital Internal Fund 214931.

## REFERENCES

[1] S. E. Halford and J. F. Marko, Nucleic acids research 32, 3040 (2004).

[2] T. Misteli, Science 291, 843 (2001).

[3] A. S. Hansen, I. Pustova, C. Cattoglio, R. Tjian, and X. Darzacq, eLife 6 (2017), 10.7554/eLife.25776.

[4] M. Ptashne, Nature 322, 697 (1986).

[5] O. G. Berg, R. B. Winter, and P. H. von Hippel, Biochemistry 20, 69296948 (1981).

[6] P. Hammar, P. Leroy, A. Mahmutovic, E. G. Marklund,O. G. Berg, and J. Elf, Science 336 (2012).

[7] J. Elf, G.-W. Li, and X. S. Xie, Science 316 (2007).

[8] M. Slutsky, M. Kardar, and L. A. Mirny, Phys. Rev. E 69, 061903 (2004).

[9] T. Hu, A. Y. Grosberg, and B. I. Shklovskii, Biophysical journal 90, 2731 (2006), arXiv:0510043 [q-bio].

[10] G.-W. Li, O. G. Berg, and J. Elf, Nature Physics 5, 294 (2009).

[11] M. Bauer and R. Metzler, Biophysical journal 102, 2321 (2012).

[12] J. Cartailler and J. Reingruber, Physical Biology 12, 046012 (2015), arXiv:1507.02452.

[13] O. Bénichou, Y. Kafri, M. Sheinman, and R. Voituriez, Phys. Rev. Lett. 103, 138102 (2009).

[14] M. A. Lomholt, B. van den Broek, S.-M. J. Kalisch,G. J. L. Wuite, and R. Metzler, Proceedings of the National Academy of Sciences of the United States of America 106, 8204 (2009).

[15] E. F. Koslover, M. A. Díaz De La Rosa, and A. J. Spakowitz, Biophysical Journal 101, 856 (2011).

[16] G. Kolesov, Z. Wunderlich, O. N. Laikova, M. S. Gelfand, and L. A. Mirny, Proceedings of the National Academy of Sciences 104, 13948 (2007), http://www.pnas.org/content/104/35/13948.full.pdf.

[17] A. Godec and R. Metzler, Phys. Rev. X 6, 041037 (2016).

[18] D. Normanno, L. Boudarène, C. Dugast-Darzacq, J. Chen, C. Richter, F. Proux, O. Bénichou, R. Voituriez, X. Darzacq, and M. Dahan, Nature communications 6, 7357 (2015).

[19] A. Amitai and D. Holcman, Phys. Rev. Lett. 110, 248105 (2013).

[20] L. Caccianini, D. Normanno, I. Izeddin, and M. Dahan, Faraday discussions 184, 393 (2015).

[21] M. Doi and S. F. Edwards, The Theory of Polymer Dynamics (Oxford: Clarendon Press, 1986).

[22] A. Amitai, A. Seeber, G. S., and D. Holcman, Cell Reports 18, 1200 (2017).

[23] O. Pulkkinen and R. Metzler, Phys. Rev. Lett. 110, 198101 (2013).

[24] A. F. Cheviakov and M. J. Ward, Math. Comput. Modelling 53, 1394 (2011).

[25] C. C. Robinett, A. Straight, G. Li, C. Willhelm, G. Sudlow, A. Murray, and A. S. Belmont, J Cell Biol 135, 1685 (1996).

[26] I. Izeddin, V. Récamier, L. Bosanac, I. I. Cissé,L. Boudarene, C. Dugast-Darzacq, F. Proux,O. Bénichou, R. Voituriez, O. Bensaude, M. Dahan, and X. Darzacq, eLife 2014, 1 (2014).

[27] M. Slutsky and L. A. Mirny, Biophysical journal 87, 4021 (2004), arXiv:0402005 [q-bio].

[28] J. Reingruber and D. Holcman, Phys. Rev.E 84, 020901 (2011).

[29] M. Sheinman and Y. Kafri, Phys. Biol. 6, 016003 (2009).

[30] O. Bénichou, C. Chevalier, B. Meyer, and R. Voituriez, Phys. Rev. Lett. 106, 038102 (2011).

[31] M. A. Lomholt, T. Ambjörnsson, and R. Metzler, Phys. Rev. Lett. 95, 260603 (2005).

[32] A. A. Shvets and A. B. Kolomeisky, The Journal of Physical Chemistry Letters 7, 5022 (2016).

[33] L. Mirny, M. Slutsky, Z. Wunderlich, A. Tafvizi, J. Leith, and A. Kosmrlj, Journal of Physics A: Mathematical and Theoretical 42, 434013 (2009).

[34] A. Amitai and D. Holcman, Physics Reports 678, 1 (2017).

[35] J. E. Lennard-Jones, Proc. R. Soc. Lond. A 106, 463477 (1924).

[36] P. G. de Gennes, Scaling Concepts in Polymer Physics (Ithaca, New York: Cornell Univ. Press, 1979).

[37] Z. Schuss, Diffusion and Stochastic Processes. An Analytical Approach (Springer-Verlag, New York, NY, 2009).

[38] C. L. Woodcock and R. P. Ghosh, Cold Spring Harbor perspectives in biology 2, a000596 (2010).

[39] S. C. Knight, L. Xie, W. Deng, B. Guglielmi, L. B. Witkowsky, L. Bosanac, E. T. Zhang, M. El Beheiry,J.-B. Masson, M. Dahan, Z. Liu, J. A. Doudna, and R. Tjian, Science 350 (2015).

[40] I. Cisse, I. Izeddin, S. Z. Causse, L. Boudarene,A. Senecal, L. Muresan, C. Dugast-darzacq, B. Hajj,M. Dahan, and X. Darzacq, Science 341, 664 (2013).

